# A genetically encoded biosensor reveals location bias of opioid drug action

**DOI:** 10.1101/254490

**Authors:** Miriam Stoeber, Damien Jullie, Toon Laeremans, Jan Steyaert, Peter W. Schiller, Aashish Manglik, Mark von Zastrow

## Abstract

Opioid receptors (ORs) precisely modulate behavior when activated by native peptide ligands but distort behaviors to produce pathology when activated by non-peptide drugs. A fundamental question is how drugs differ from peptides in their actions on target neurons. One way that drugs can differ is by imposing selective effects on the conformational equilibrium of individual ORs. We wondered if drugs can also impose selective effects on the location of OR activation in individual OR-expressing neurons. Here we develop a genetically encoded biosensor that directly localizes ligand-induced activation and deactivation of ORs in living cells, and use it to generate the first real-time map of the spatiotemporal organization of μ‐ and δ-OR activation in neurons. Peptide agonists produce a characteristic activation pattern initiated in the plasma membrane and propagating to endosomes after receptor internalization. Drugs produce a different activation pattern by uniquely driving OR activation in the somatic Golgi apparatus and extending throughout the dendritic arbor in Golgi outposts. These results demonstrate a new approach to probe the cellular basis of neuromodulation and reveal that drugs profoundly distort the spatiotemporal landscape of neuronal OR activation.

## Introduction

Opiate alkaloid drugs such as morphine are among the most effective agents known for alleviating pain. However, such drugs produce significant toxicity and have high abuse potential. These factors have contributed to opioid addiction becoming a large and growing public health problem globally, which has now reached epidemic proportions in the US (Volkow and McLellan, 2016). Opiate drugs produce their biological effects by binding to the same subfamily of G protein-coupled receptors (GPCRs), the opioid receptors (ORs), as endogenously produced opioid peptide ligands (Kieffer and Evans, 2008). This supports a long-held view (Bradbury et al., 1976) that opiate drugs mimic the actions of peptide ligands at the level of individual receptor-expressing target neurons. Despite an urgent need to develop improved analgesic therapies, and compounded by the fact that opiate drugs remain mainstays in the pharmacological management of severe pain, it remains unclear to what degree opiate drugs differ in their cellular effects (Thompson et al., 2015). It also remains unknown if opiate drugs, beyond producing cellular effects that are similar to peptide ligands, have the potential to produce discrete or additional receptor-based effect(s) that opioid peptides cannot.

Drugs are well known to differ substantially from peptides in physicochemical properties that affect membrane permeability and drug bioavailability within tissues (Weber et al., 1993). However, such differences have been thought to have little impact at the cellular level because present models assume that receptor activation is restricted to the plasma membrane (PM) (Zastrow and Williams, 2012). It has become clear recently that a number of other GPCRs are subject to ligand-dependent activation at internal membrane locations as well as the PM (Calebiro et al., 2010; Irannejad et al., 2015; Jong et al., 2017). With this in mind, and in light of the exquisitely high degree of membrane compartmentation that is characteristic of neurons, we wondered if ORs have the ability to undergo ligand-dependent activation at more than one membrane location in neurons. If so, might chemically distinct ligands differ in the subcellular location(s) at which they produce OR activation?

Resolving the precise subcellular location(s) at which wild type ORs are activated by ligands was previously impossible due to a lack of suitable experimental tools. Recent advances in GPCR structural biology have made feasible the development of a new class of genetically encoded probes, called conformational biosensors, which detect activated receptors based on a defined activation-associated conformational change (Manglik et al., 2017). Conformational biosensors have revealed that catecholamine receptors undergo ligand-induced activation both in the PM and at internal membrane sites (Irannejad et al., 2017; 2013). No such biosensor has yet been described for detecting ligand-dependent activation of ORs or any peptidergic GPCR. Further, no previous study has investigated the ability of conformational biosensor technology to detect activation of any receptor in neurons.

Here we describe a conformational biosensor that specifically senses ligand-induced activation of μ‐ and δ-ORs in living neurons. We demonstrate that this tool provides precise spatial and temporal resolution of OR activation and deactivation *in situ* and with minimal perturbation of function. Using this novel genetically encoded probe, we show that neuronal OR activation is not restricted to the cell surface as previously assumed. Instead, activation begins in the PM and propagates to endosomes after ligand-induced OR internalization. We then demonstrate that both antagonist and agonist drugs distort this pattern. In particular, non-peptide drugs, including the prototypic opiate alkaloid morphine, drive a discrete wave of internal OR activation in the somatic Golgi apparatus and distributed throughout the dendritic arbor in Golgi outposts. These results reveal a characteristic pattern of subcellular OR activation generated by peptides and its profound distortion by drugs. We propose a new principle of biased drug action that is imposed at the level of individual target neurons and manifest through the spatiotemporal landscape of receptor activation.

## Results

### A conformational biosensor for direct and specific detection of OR activation

To develop an OR activation biosensor we began with clones selected from a camelid-derived antibody fragment (nanobody) library that bind *in vitro* to μ-OR specifically in an active (agonist-bound) relative to inactive (antagonist-bound) conformation (Figure 1A) (Huang et al., 2015; Manglik et al., 2012). We selected four nanobody clones in which key residues that engage μ-OR in the active structure are conserved (Figure S1A). We prepared N‐ and C-terminal fluorescent protein fusions from each nanobody clone and tested them for cytoplasmic expression. The ability of nanobody fusion proteins to act as sensors of active-conformation ORs was first examined by co-expression with μ-OR in human embryonal kidney cells and imaging of cells using total internal reflection fluorescence microscopy (TIR-FM). TIR-FM allowed studying recruitment to the PM because it selectively detects fluorophores at the cell surface (Figure 1B). Several constructs exhibited ligand-dependent recruitment to the PM and N-terminally tagged Nb33 (named OR-sensor) was selected for further study. In cells not exposed to opioid ligands, little fluorescence of OR-sensor was observed in the TIR-FM illumination volume, consistent with localization in the cytoplasm. Activation of co-expressed μ-OR with a peptide agonist (DAMGO) triggered a pronounced increase of OR-sensor fluorescence in the TIR-FM field (Figures 1C and 1D, Movie S1). PM recruitment of OR-sensor had a rapid onset (t_1/2_ <20 s) after DAMGO application and was rapidly reversed (t_1/2_ ∼20 s) after agonist washout or addition of the competitive antagonist naloxone (Figure 1E).

**Figure 1:**
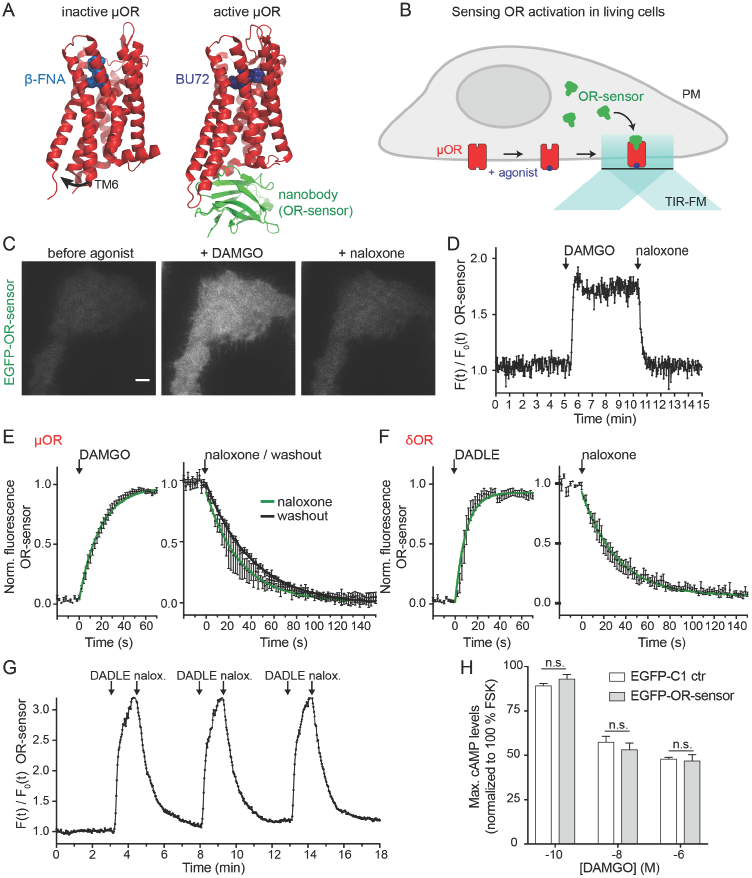
A conformational biosensor for direct detection of OR activation. **(A)** Crystal structures of inactive, ß-FNA-bound (4DKL) and active, BU27-bound (5C1M) μ-OR (red). The active conformation is stabilized by a nanobody (Nb39, green), named OR-sensor. Ligands are in blue. Arrow indicates outward movement of TM6 upon activation. See also Figure S1. **(B)** Schematic of OR-sensor and μ-OR localization in living cells and expected OR-sensor re-localization upon agonist addition. Total internal reflection fluorescence microscopy (TIR-FM) light beam is indicated. PM = plasma membrane. **(C)** TIR-FM images of a time series of a HEK293 cell, expressing EGFP-OR-sensor (and μ-OR, not shown). Medium changes to DAMGO (1 μM, agonist) or Naloxone (10 μM, antagonist) by perfusion. Scale bar: 10 μm. **(D)** EGFP-OR-sensor fluorescence intensity during TIR-FM time-lapse series with media changes to DAMGO and Naloxone. 2 s between frames. F_0_ = average fluorescence intensity of first 15 frames. See also Figure S2. **(E & F)** Recruitment and detachment kinetics of OR-sensor to μ-OR or δ-OR in the PM, measured by EGFP-OR-sensor fluorescence intensity increase using TIR-FM. 2 s between frames. Normalization of EGFP-intensity values (range [0-1]). Data fit with one-phase exponential association or one-phase decay formula. **(E)** HEK293 cells expressing μ-OR and OR-sensor. Left: DAMGO (1 μM) was applied by media perfusion starting at t = 0. (n=9, average +/− sem). Right: medium switch from DAMGO (1 μM) to naloxone (10 μM) or no agonist (washout) (naloxone: n=4, washout n= 8, average +/− sem). **(F)** HEK293 cells expressing δ-OR and OR-sensor. Left: DADLE (1 μM) was applied by media perfusion starting at t = 0 (n=5, average +/− sem). Right: Medium switch from DADLE (1 μM) to naloxone (10 μM) (n=5, average +/− sem). **(G)** EGFP-OR-sensor fluorescence intensity measured by TIR-FM in a HEK293 cell expressing OR-sensor and δ-OR (not shown). Repeated media switches to 1 μM DADLE (1 min), 10 μM naloxone (2 min), and wash out (2 min). F_0_ = average fluorescence intensity of first 15 frames. **(H)** Maximal cAMP response measured by pGloSensor cAMP sensor in HEK293 cells stably expressing μ-OR and transiently expressing EGFP-control or EGFP-OR-sensor. Cells were stimulated with Forskolin (FSK, 2 μM) and different concentrations of DAMGO. Data normalized to 100% FSK control. (n=3, average +/− SD, n.s. = no significant difference, unpaired t-test).

The δ-opioid receptor (δ-OR) is closely homologous in structure to μ-OR, and all residues contacting OR-sensor in the activated conformation of μ-OR are conserved in δ-OR (Figure S1B). Verifying that OR-sensor also detects ligand-activated δ-OR, the peptide agonist DADLE triggered rapid OR-sensor recruitment to the PM that was rapidly reversible by antagonist (Figure 1F). Verifying receptor-dependence of this response, neither DAMGO nor DADLE produced a recruitment signal in cells not expressing ORs (Figure S2). Verifying receptor-specificity, expression of the Gi-coupled M2 muscarinic receptor at similar levels as OR failed to produce detectable recruitment of OR-sensor following application of the muscarinic agonist carbachol (Figure S2).

A potential caveat of conformational biosensors is that, because the core nanobody structure binds directly to the active receptor conformation, they could ‘force’ receptor activation in a ligand-independent manner. This is unlikely because OR-sensor recruitment to the PM was not observed in the absence of agonist, and because recruitment of OR-sensor was rapidly reversible upon agonist removal. Emphasizing reversibility of OR-sensor recruitment, alternating agonist and antagonist application to the same cell produced repeated rounds of OR-sensor recruitment and dissociation from the PM (Figure 1G). Another potential caveat is that OR-sensor could block receptor function. This was also not the case because OR-sensor did not produce any detectable effect on the ability of receptors to mediate ligand-dependent inhibition of adenylyl cyclase, an assay requiring receptor coupling to Gi protein (Figure 1H). Together, these results indicate that OR-sensor provides direct and specific detection of OR activation and deactivation in living cells with minimal perturbation of receptor function.

### Peptide agonists drive sequential OR activation waves in the PM and endosomes

Peptide-induced activation of both μ-OR and δ-OR promotes rapid, clathrin-dependent endocytosis of receptors. Peptide binding to GPCRs can be reduced by acidification and proteolysis (Grady et al., 1995; Gupta et al., 2014), and current models of OR regulation involving endocytosis assume that OR activation is restricted to the PM (Zastrow and Williams, 2012). The development of OR-sensor provided an opportunity to directly test this assumption. We labeled FLAG-tagged μ-OR or δ-OR in the cell surface with a fluorescent monoclonal antibody and used confocal microscopy to simultaneously image receptor and OR-sensor in living cells. μ-OR remained in the PM in the absence of agonist and accumulated in endosomes within several minutes after application of DAMGO (Figure 2A, Movie S2). OR-sensor was transiently recruited to the PM, consistent with the TIR-FM observations, and then localized to OR-containing endosomes (Figure 2A). Similar results were observed in experiments using δ-OR rather than μ-OR (Figure 2B, Movie S3). The kinetics of receptor and OR-sensor recruitment to endosomes were estimated by three-color confocal imaging, using Early Endosome Antigen 1 (EEA1) as a marker of the early endosome population (Figure 2C). We measured the average intensity of μ-OR and OR-sensor fluorescence within a mask defined by EEA1 as a function of time after peptide agonist addition. Accumulation of μ-OR in endosomes began ∼1 min after DAMGO application and reached a plateau at ∼20 min, consistent with previously published rates of μ-OR endocytosis and approach to steady state through continuous endocytosis and recycling that occurs in the presence of peptide agonist (Henry et al., 2012). Ligand-induced OR-sensor recruitment to endosomes lagged receptor accumulation in this compartment and also reached a plateau within ∼20 min (Figure 2D). These results indicate that active-conformation ORs are not restricted to the PM but are also present in endosomes.

**Figure 2:**
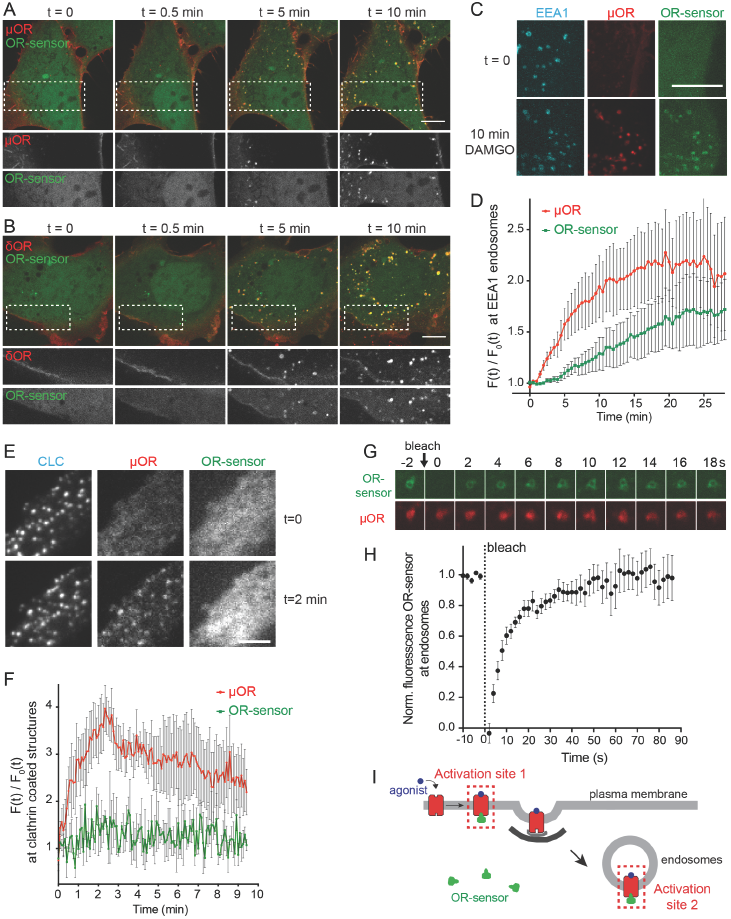
An endosome-localized wave of OR activation. Confocal images of time series of HEK293 cells, expressing EGFP-OR-sensor and FLAG-μ-OR **(A)** or FLAG-δ-OR **(B).** Receptors were surface-labeled with anti-FLAG M1-AF555. 10 μM DAMGO (A) or DADLE (B) was added at t=0. Boxed areas are displayed separately for both fluorophores below respective images. Scale bar: 10 μm. **(C)** Maximum Z-projection of 12 confocal slices of a HEK293 cell, expressing EGFP-OR-sensor, FLAG-μ-OR (surface-labeled with M1-AF647, pseudo-colored red), and an early endosomal marker (DsRed2-EEA1-FYVE, pseudo-colored in blue) before (t=0) and after (t=15 min) adding DAMGO (10 μM). Images are representative examples used for quantification shown in (D). Scale bar 10 μm. **(D)** Quantification and kinetics of μ-OR and OR-sensor recruitment to endosomes. EEA1-positive vacuoles were used as quantification mask. FLAG-μ-OR and EGFP-OR-sensor fluorescence intensity was measured within the mask at each time point of the image series. F_0_ = average fluorescence signal in EEA1 mask before agonist addition. Movie length 25 min, 30 s between Z-stacks (n=5, average +/− sem). **(E)** TIR-FM images of a time series of a HEK293 cell, coexpressing clathrin-light-chain (CLC)-DsRed, FLAG-μ-OR (surface-labeled with M1-AF647) and EGFP-OR-sensor. DAMGO (10 μM) was added at t=0. Scale bar 5 μm. See also Figure S3. **(F)** Quantification of μ-OR and OR-sensor intensity in clathrin-coated structures vs. PM over time. CLC-positive spots were used as quantification mask. F_0_ = average fluorescence signal in CCPs before agonist addition. Movie length 10 min, 5 s between frames (n=3, average +/− sem). **(G)** Fluorescence-recovery after photobleaching (FRAP) series displaying intensity of EGFP-OR-sensor (photobleached) and μ-OR (surface-labeled with M1-AF647) at a bleached endosome, 2 s intervals. **(H)** FRAP of EGFP-OR-sensor at μ-OR-loaded endosomes in HEK293 cells, stimulated for 15 min with DAMGO (10 μM). Normalized EGFP-OR-sensor florescence in bleached area over time. 2 s interval between frames during acquisition (n=7, average +/− sem) **(I)** Scheme depicting the two distinct sites of OR-sensor recruitment to peptide ligand-activated receptors: activation site 1: plasma membrane, activation site 2: endosomes.

A potential caveat to this interpretation is that OR-sensor could accumulate in endosomes as an artifact of a persistent receptor-biosensor complex that forms at the PM and remains associated during endocytosis. Two lines of evidence argue against this. First, TIR-FM time-lapse analysis showed that ORs robustly cluster in clathrin-coated pits (CCPs) without concomitant recruitment of OR-sensor to the endocytic site (Figures 2E and 2F), and CCPs pinch off from the PM containing ORs but not associated with OR-sensor (Figure S3). Second, when we applied fluorescence recovery after photobleaching (FRAP) analysis to examine exchange kinetics of OR-sensor at individual μ-OR-containing endosomes, the OR-sensor signal fully recovered with a half-time of ∼15 sec (Figures 2G and 2H), much faster than the kinetics of receptor trafficking (Figure 2D). This rules out the possibility that OR-sensor localization to endosomes occurs as an artifact of stable receptor-biosensor complex formation. To the contrary, OR-sensor exchanges remarkably rapidly and therefore provides a continuous readout of the receptor’s activation state. Together, these results reveal spatiotemporal organization of cellular OR activation induced by peptide agonists, with sequential activation occurring in the PM and then in endosomes following the process of ligand-induced internalization of ORs (Figure 2I).

### Endosome OR activation is ligand-dependent but differs from the PM wave in duration and antagonist regulation

We next tested if OR activation in endosomes is indeed dependent on the presence of agonist ligand. If endosomal activation requires agonist binding to receptors, as it does at the PM, we expected competitive antagonist ligands to reverse endosomal OR-sensor recruitment depending on the ability of the antagonist to access internalized receptors (Figure 3A). Activation of μ-OR in endosomes remained ligand-dependent because naloxone, a membranepermeant alkaloid antagonist rapidly and fully reversed OR-sensor localization to endosomes within 30 s, a time period shorter than required for μ-OR recycling. Verifying this, internalized receptors were still present in endosomes after OR-sensor dissociation (Figure 3B). Interestingly, while naloxone reversed the OR-sensor recruitment signal in <30 s, full reversal of the activation signal after agonist removal from the culture medium required >30 min. Slow reversal was also observed in the presence of a peptide antagonist (CTOP) that is membraneimpermeant (Figure 3C). Similar results were obtained in experiments investigating endosomal OR-sensor recruitment mediated by δ-OR (Figure 3D). These results indicate that OR activation in endosomes is ligand-dependent but differs from PM activation in its resistance to washout and differential access to antagonists.

**Figure 3:**
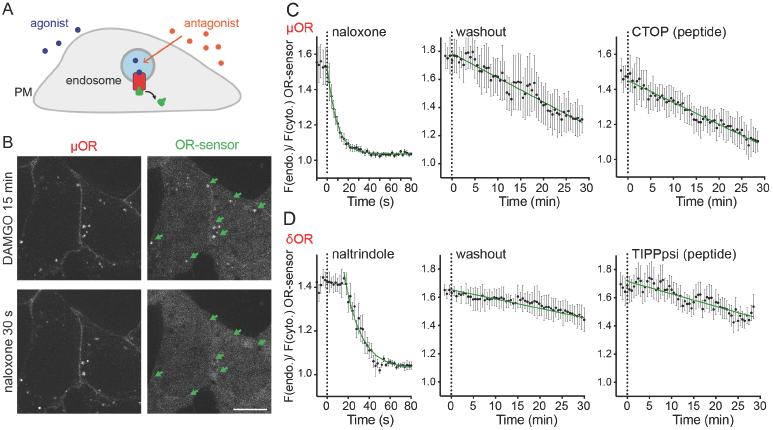
Endosomal OR activation is ligand-dependent. **(A)** Detecting OR activation in endosomes using OR-sensor. **(B)** Confocal images of a HEK293 cell expressing μ-OR (surface-labeled with M1-AF555) and EGFP-OR-sensor. Top: 15 min after adding DAMGO (1 μM). Bottom: 30 s after adding Naloxone (10 μM). Scale bar: 10 μm. **(C)** Quantification of EGFP-OR-sensor intensity at μ-OR-containing endosomes (treated for 15 min with 1 μM DAMGO) upon agonist washout or after adding 10 μM of non-peptide (naloxone) or peptide (CTOP) antagonist. EEA1-endosomes were used as quantification mask and intensity normalized to cytosolic OR-sensor signal at each time point. Naloxone: 2 s between frames, n=6, washout: 30 s between frames, n=3, CTOP: 30 s between frames, n=4, average +/− sem. **(D)** Quantification of EGFP-OR-sensor intensity at δ-OR-containing endosomes (15 min 1 μM DADLE) upon agonist washout or after adding 10 μM of non-peptide (naltrindole) or peptide (TIPPpsi) antagonist. δ-OR-positive endosomes were used as quantification mask and intensity normalized to cytosolic signal of OR-sensor at each time point. Naltrindole: 2 s between frames, n=5, washout: 30 s between frames, n=5, TIPPpsi: 30 s between frames n=5, average +/− sem.

### Sequential waves of OR activation in neurons

We next investigated whether OR-sensor can detect receptor activation in neurons, and chose primary cultures of striatal medium spiny neurons that are known to express μ-OR and δ-OR endogenously in a fraction of neurons. To start we expressed OR-sensor together with epitopetagged OR to allow reliable detection of both proteins in the same cells, and carried out simultaneous imaging of both using time-lapse confocal microscopy. Prior to agonist addition, OR-sensor showed a diffusive cytosolic distribution in soma and dendrites (Figure 4A). DAMGO addition caused a rapid accumulation of OR-sensor at the PM, which was particularly evident in dendritic processes (Figure 4B, left and middle panels). Within minutes thereafter, DAMGO addition induced μ-OR redistribution from the PM to endosomes that appeared as punctate structures in soma and dendrites (Figures 4A and 4B, Movie S4). Strikingly, the μ-OR-loaded endosomes also recruited OR-sensor. We verified that these structures indeed represent endosomes by co-labeling with EEA1 (Figure 4C) and then used EEA1 as marker for quantifying μ-OR and OR-sensor recruitment to endosomes over time. As in non-neural cells, OR-sensor recruitment to the endosome compartment was strong and appeared to lag μ-OR accumulation (Figure 4D). Further, the biosensor signal at endosomes was reversed within seconds after addition of naloxone, a time point clearly preceding μ-OR exit from endosomes (Figure 4E).

**Figure 4:**
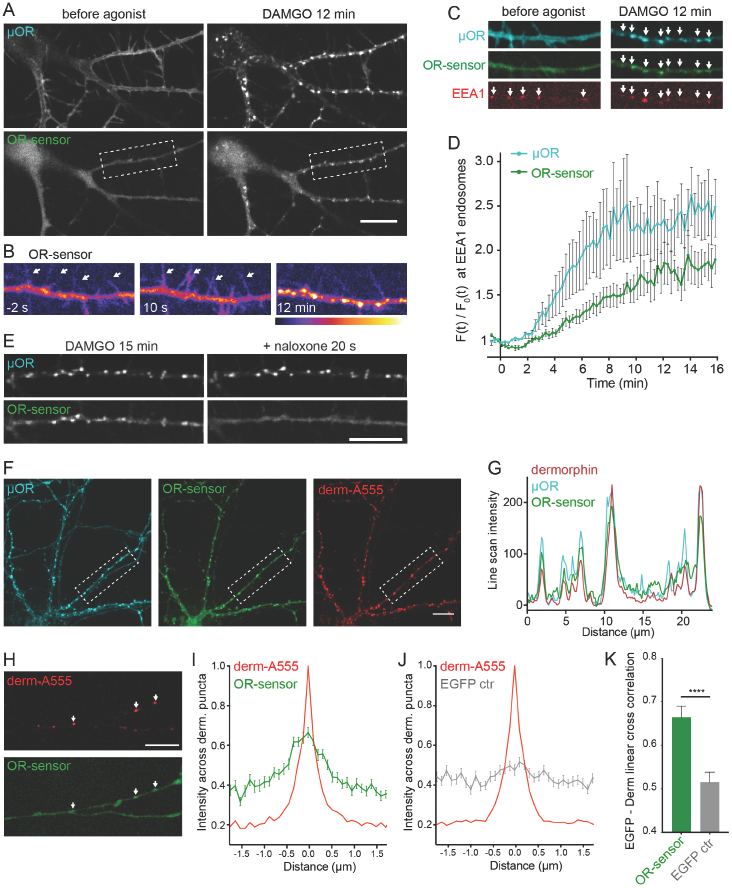
OR activation in somatodendritic endosomes. **(A)** Striatal neuron (12 DIV), expressing FLAG-μ-OR (surface-labeled with M1-AF555) and EGFP-OR-sensor. Images of a time series, before and 12 min after adding DAMGO (10 μM). Scale bar: 10 μm. **(B)** Pseudocolor images (low to high intensity) of OR sensor, corresponding to boxed area in (A). Dendrite before (-2 s), immediately after (10 s) and 12 min after adding DAMGO. Arrows depict dendritic processes. **(C)** Dendrite of a striatal neuron (12 DVI), expressing FLAG-μ-OR (surface-labeled with M1-AF647), OR-sensor, and EEA1 before agonist and 12 min after adding DAMGO (10 μM). Arrows depict EEA1-positive endosomes that contain μ-OR and recruit OR-sensor after agonist addition. **(D)** Quantification and kinetics of μ-OR and OR-sensor recruitment to endosomes in dendrites. EEA1-positive vacuoles were used as quantification mask. F_0_ = average fluorescence signal before agonist addition. Movie length 16 min, 5 s between frames (n=5, average +/− sem). **(E)** Localization of μ-OR and OR-sensor in a dendrite 15 min after adding DAMGO, and 20 s after adding naloxone (10 μM). **(F)** Confocal images showing localization of FLAG-μ-OR (surface-labeled with M1-AF647, blue), EGFP-OR-sensor, and fluorescent dermorphin (derm-A555, red) in striatal neurons 15 min after derm-A555 addition. **(G)** Fluorescence intensity profile (line scan) of dermorphin (red), μ-OR (blue), and OR-sensor along dendritic process, boxed in (F). **(H)** Dendrite of a striatal neuron (14 DVI) expressing OR-sensor, incubated with derm-A555 (1 μM) for 15 min. Arrows depict dermorphinloaded vacuoles that co-localize with OR-sensor. **(I,J)** Average fluorescence intensity profile (line scans) of EGFP-OR-sensor (I) or EGFP control (J) along dendrite across derm-A555-labeled endosomes in striatal neurons. (OR-sensor: n=110 endosomes from 15 cells, EGFP-control: n=102 endosomes from 18 cells, average +/− sem.) **(K)** Peak cross correlation at derm-A555 puncta. EGFP-OR-sensor = 0.6645 ± 0.0257, EGFP control = 0.5152 ± 0.0233, ****p <0.0001, unpaired t-test.

Our antagonist and washout experiments had shown that the active conformation of ORs at the PM depended on the presence of agonist. To verify this, we asked if μ-OR-containing endosomes that recruit OR-sensor also contain agonist peptide. We used the opioid peptide dermorphin conjugated to AlexaFluor555 (derm-A555), shown previously to retain biological activity (Arttamangkul et al., 2000). Verifying this, dermorphin application to neurons drove the rapid internalization of μ-OR as well as OR-sensor recruitment to μ-OR-containing endosomes (Figure 4F, left and middle panel). Importantly, all of the endosomes containing activated μ-OR colocalized with dermorphin (Figures 4F and 4G), demonstrating that opioid peptide is indeed present at sites of endosomal μ-OR activation.

Because fluorescent dermorphin is sufficiently sensitive to label endosomes containing endogenous μ-ORs (Arttamangkul et al., 2006), we next asked if endosomal activation also occurs in neurons expressing only native receptors. Indeed, dermorphin-containing endosomes were detected throughout dendrites in a subset of neurons and, at many of these endosomes, OR-sensor recruitment was visually evident (Figure 4H). We used cross-correlation analysis to assess endosomal OR-sensor recruitment mediated by endogenous ORs in an unbiased manner and across hundreds of endosomes in multiple experiments. This analysis revealed a clear cross-correlation signal between OR-sensor fluorescence intensities and endosomes marked by derm-A555 (Figure 4I). In contrast, no cross-correlation signal was detected in control experiments using soluble EGFP in place of EGFP-OR-sensor (Figures 4J and 4K). These results indicate that OR-sensor can detect activation of ORs expressed at native levels and that opioid peptide can drive endosomal activation of endogenous ORs in neurons.

### Opioid drugs distort the spatiotemporal pattern of cellular OR activation

Another unexpected result was obtained when we examined the effect of agonist drugs. The alkaloid agonist etorphine drove OR-sensor recruitment to a perinuclear membrane compartment within seconds after application, before detectable internalization and almost simultaneous with OR activation in the PM (Figure 5A, Movie S5). Morphine produced a similar effect (Figure 5B). The compartment to which drugs drove OR-sensor recruitment did not originate from the endocytic pathway because it failed to label with receptors internalized from the PM as visualized using our surface labeling protocol (Figure 5A). Neurons are known to contain an internal membrane pool of ORs that originates from the biosynthetic rather than endocytic pathway and localizes at steady state to ER and Golgi membranes (Erbs et al., 2014; Scherrer et al., 2006). Consistent with this, an internal pool of μ‐ and δ-ORs was detected by immunostaining of fixed cells after permeabilization, and this pool colocalized with the Golgi marker Giantin (Figure 5C, Figure S4A). We used GFP-fused OR constructs to visualize this internal membrane pool in living cells and observed that alkaloid agonists drove rapid OR-sensor recruitment precisely to this compartment (Figure 5D, Movie 6). Verifying specificity of the activation signal, OR-sensor recruitment to Golgi membranes was rapidly reversed by opiate antagonist (Figure 5D).

**Figure 5:**
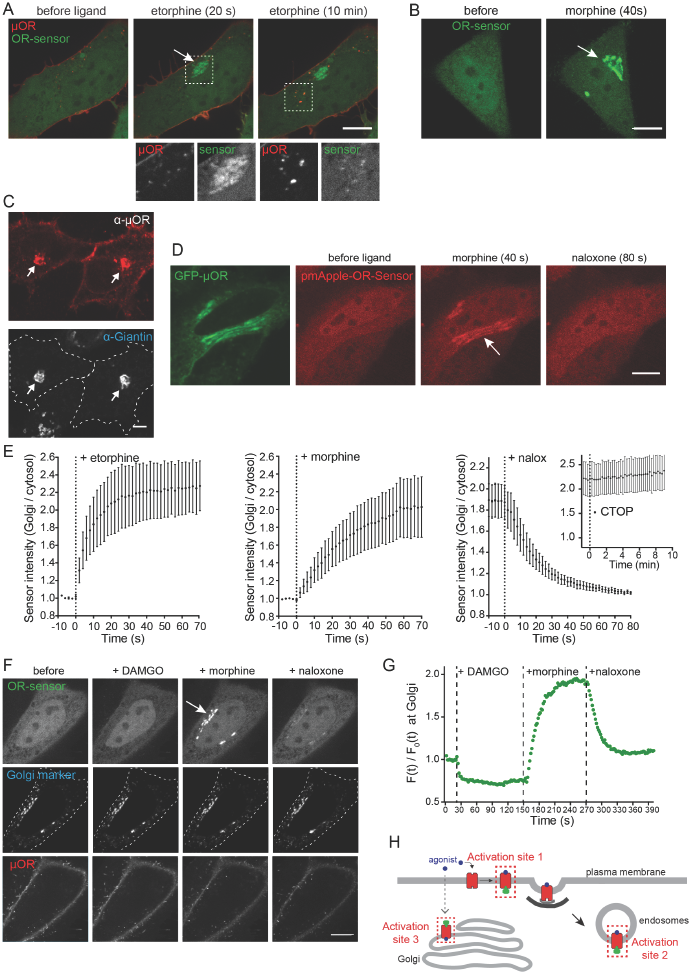
Non-peptide drugs activate Golgi-localized ORs. **(A)** Confocal images of a time series of a HEK293 cell, expressing EGFP-OR-sensor and FLAG-μ-OR (surface-labeled with anti-FLAG M1-AF555). Etorphine (1 μM) was added at t=0. Boxed areas are displayed separately for both fluorophores below respective images. **(B)** Confocal images of time series of a HeLa cells, expressing EGFP-OR-sensor and FLAG-μ-OR (not depicted). OR-sensor localization is shown before and 20 s after adding morphine (1 μM). **(C)** Confocal images of HeLa cells, expressing FLAG-μ-OR. Cells were fixed, permeabilized, and immunolabelled with anti-FLAG (red) and anti-Giantin (grey) antibodies. The internal OR pool co-localizes with Golgi marker (arrows). See also Figure S4A. **(D)** Rapid activation of Golgi-localized μ-ORs by morphine. Confocal images of a time series of a HeLa cell, expressing μ-OR-EGFP and mCherry-OR-sensor. Cell was treated with morphine (1 μM), followed by naloxone (10 μM). **(E)** Quantification and kinetics of EGFP-OR-sensor intensity at Golgi apparatus upon agonist or antagonist addition in HeLa cells, expressing OR-sensor, GalT-DsRed, and μ-OR. GalT-marked Golgi apparatus was used as quantification mask and intensity normalized to cytosolic OR-sensor signal at each time point. Morphine n=4, etorphine n=3, 1 μM etorphine followed by 10 μM naloxone n=6, or by 10 μM CTOP n=5. 2 s intervals for non-peptide ligands, 15 s intervals for peptide ligand (CTOP), average +/− sem. See also Figure S4B and S4C. **(F)** Confocal images of a time series of a HeLa cell, expressing EGFP-OR-sensor, GalT-DsRed (Golgi-marker), and FLAG-μ-OR (surface labeled with M1-647). Cell was treated with DAMGO (10 μM, t = 20 s), morphine (1 μM, t = 150 s), and naloxone (10 μM, t = 270 s). **(G)** Quantification of EGFP-OR-sensor intensity at Golgi throughout time series shown in (F). 10 s between frames. **(H)** Scheme depicting three distinct sites of OR-sensor recruitment to activated receptors following etorphine addition: activation site 1: plasma membrane, activation site 2: endosomes, activation site 3: Golgi apparatus. All scale bars: 10 μm.

To quantify the kinetics of the Golgi-localized activation signal, we labeled the Golgi apparatus in living cells with GalT-DsRed and quantified OR-sensor recruitment throughout confocal time series after application of agonist drug. Both morphine and etorphine drove robust recruitment of OR-sensor to Golgi-localized μ-OR with t_1/2_ < 20s (Figure 5E). Etorphine, as well as the distinct non-peptide δ-OR agonist ARM390, produced a similarly rapid and robust Golgi activation response in cells expressing δ-OR (Figure S4B and S4C). All of these non-peptide drugs are thought to be relatively membrane-permeant when compared to peptides, and the remarkably fast kinetics of the Golgi-localized OR activation response argues that ligand access from the cell surface is not likely mediated by membrane trafficking because this process typically has slower kinetics. Instead, it suggested that non-peptide drugs selectively produce Golgi-localized OR activation because they diffuse across membranes. Supporting this hypothesis, Golgi-localized activation of μ-OR and δ-OR was reversed rapidly after application of naloxone and naltrindole, membrane-permeant alkaloid antagonists, but it persisted after application of CTOP, TIPPpsi, or ICI-174,864 that are peptide-based antagonists thought to penetrate membranes slowly (Figure 5E, Figure S4C).

To directly verify that Golgi-localized OR activation is specific to drugs, we challenged individual μ-OR-expressing cells sequentially – first with the opioid peptide agonist DAMGO and then with the alkaloid agonist morphine – while recording time-lapse series of OR-sensor localization (Figure 5F). Strikingly, DAMGO failed to produce detectable recruitment of OR-sensor to Golgi membranes but subsequent application of morphine produced a robust activation signal that was receptor-dependent and reversible by naloxone (Figure 5F and 5G). Together these results identify Golgi membranes as a discrete internal site of OR activation that is rapidly accessible to alkaloid (and other non-peptide) drugs but is inaccessible to opioid peptides (Figure 5H).

### OR activation in somatic Golgi and Golgi outposts of neurons

Striatal neurons also contained a Golgi-localized pool of ORs, as detected in the soma by colocalization with GalT (Figure 6A). Non-peptide agonists including etorphine (Figure S5A) and morphine (Figure 6B, Movie S7) drove rapid recruitment of OR-sensor to this internal membrane pool. We verified that the Golgi pool is not accessible to ORs internalized from the PM (Figure S5B). Moreover, we detected rapid kinetics of drug-induced OR activation in Golgi membranes of neurons and its reversibility by membrane-permeant antagonist (Figure 6C). Therefore, non-peptide drugs rapidly and specifically promote OR activation in the somatic Golgi apparatus of neurons.

**Figure 6:**
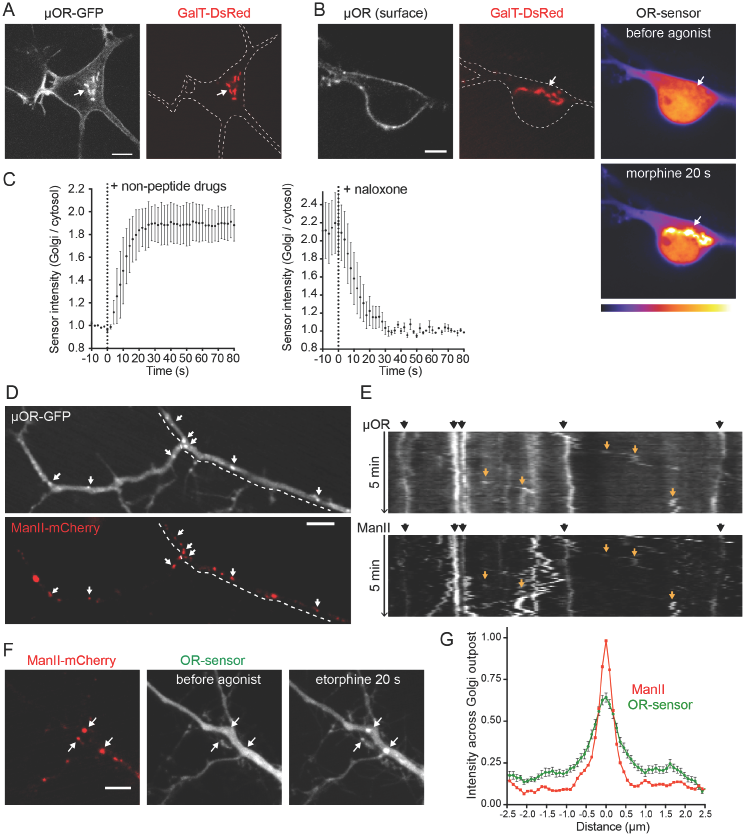
OR activation in somatic Golgi and Golgi outposts of neurons. **(A)** Soma of a striatal neuron (12 DIV), expressing μ-OR-EGFP and GalT-DsRed2 (Golgimarker). Internal μ-ORs co-localize with the Golgi marker (arrow). **(B)** Soma of striatal neuron (14 DIV), expressing FLAG-μ-OR (surface-labeled with M1-AF647), GalT-DsRed2, and OR-sensor (pseudocolored low to high intensity). OR-sensor distribution is depicted before agonist and 20 s after adding morphine (1 μM). See also Figure S5. **(C)** Quantification and kinetics of EGFP-OR-sensor intensity at somatic Golgi upon non-peptide agonist or antagonist addition in striatal neurons, expressing OR-sensor, FLAG-μ-OR, and GalT-DsRed. GalT-marked Golgi was used as quantification mask. Time series with 5 s intervals. Left: Averaged data using non-peptide drugs, from n=2 morphine (1 μM) and n=2 etorphine (1 μM). Left: 1 μM etorphine followed by naloxone (10 μM), n=3. Average +/− sem. **(D)** Dendrite of a striatal neuron (13 DIV), expressing μ-OR-EGFP and ManII-mCherry (Golgi-outpost (GOP) marker). Puncta of μ-ORs co-localize with GOP (arrows). **(E)** 5 min kymograph along the dendritic branch highlighted by dashed line in (D). Black arrows (top): stable dendritic GOPs appear as straight, vertical lines. Orange arrows (in kymograph): sloped traces of mobile secretory carriers. **(F)** Dendrite of a striatal neuron (12 DIV), expressing ManII-mCherry, EGFP-OR-sensor, and FLAG-μ-OR (not depicted). OR-sensor distribution before agonist and 20 s after adding etorphine (1 μM). GOPs recruit OR-sensor upon receptor activation (arrows). **(G)** Average fluorescence intensity profile (line scan) of EGFP-OR-sensor across ManII-labeled GOPs after adding etorphine. n=47 GOPs from 14 cells, average +/− sem. All scale bars: 10 μm.

In addition to the classical Golgi apparatus localized in the soma, many neurons contain functional Golgi-related membrane clusters in dendrites that are called Golgi outposts (Horton and Ehlers, 2003; Ori-McKenney et al., 2012; Pierce et al., 2001). We verified the presence of such membrane structures, marked by Alpha-mannosidase 2 (ManII), throughout dendrites of medium spiny neurons and established that they contain ORs (Figure 6D). Consistent with the characteristic properties of Golgi outposts (Horton and Ehlers, 2003), these structures were frequently observed at dendritic branch points and were immobile in live image series (Figure 6E, black arrows). We also observed mobile OR-containing membrane structures that appeared to emerge from static ManII-marked clusters (Figure 6E, orange arrows), consistent with transport carriers derived from Golgi outposts. Together, these observations raised the question of whether non-peptide opiate drugs, in addition to producing a discrete component of OR activation in the somatic Golgi apparatus, might have the ability to drive additional internal membrane activation more broadly by accessing Golgi outposts. This was indeed the case because alkaloid agonists produced robust OR-sensor recruitment to ManII-marked membrane structures throughout dendrites (Figure 6F and 6G). Like Golgi activation in the soma, OR-sensor recruitment to dendritic Golgi outposts occurred in <20 sec, distinguishing this component of OR activation temporally from OR activation in endosomes that develops over several minutes (Figure S5). Taken together, these results indicate that the ability of opiate alkaloids to drive a rapid and discrete component of OR activation in Golgi membranes is not restricted to the soma. Rather, OR activation by drugs extends throughout the dendritic arbor via Golgi outposts.

## Discussion

The present study establishes a genetically encoded conformational biosensor that is capable of directly resolving the subcellular location of OR activation in living neurons and in real time. To our knowledge, the present results are the first to describe a nanobody-based conformational biosensor for detecting activation of any peptidergic GPCR, and the first to apply conformational biosensor technology to assess any receptor type in neurons. Our results delineate a characteristic spatiotemporal pattern of neuronal OR activation that is produced by opioid peptides and reveal distinct activation effects of opioid drugs, thereby suggesting a new cellular framework for understanding specificity and diversity of opioid drug action at the level of individual target neurons.

The traditional understanding of OR activation in neurons holds that ligand-dependent activation of receptors is restricted to the cell surface. While neuropeptides, including opioids, are clearly influenced by endosome acidification and proteases (Grady et al., 1995; Gupta et al., 2014), the idea that ORs are necessarily inactive in endosomes is based on assumption. The development of OR-sensor provided an opportunity to test this assumption directly. Unexpectedly, we found endosomes to represent a discrete site of ligand-dependent OR activation. Using OR-sensor, we delineated a characteristic spatiotemporal pattern of OR activation that is elicited by peptide ligands in living neurons, beginning in the PM and propagating to endosomes after receptors undergo ligand-induced internalization.

Endosomal OR activation remains dynamically ligand-dependent but differs from activation in the PM due to ligand trapping. Accordingly, endosomal OR activation is longer-lasting than activation in the PM, and we directly demonstrate slow reversal after peptide removal from the extracellular milieu. This suggests that OR activation in endosomes provides a form of cellular ‘memory’ of previous opioid exposure. The effect of membrane-permeant antagonists such as naloxone is different from agonist washout because naloxone rapidly reverses endosomal OR activation. This suggests that administration of an opiate antagonist drug does not simply mimic the effect of agonist removal but, instead, it fundamentally distorts the neuronal OR activation landscape by blocking or reversing a discrete component that would normally persist. Alkaloid agonists distort the activation landscape even more profoundly by driving conformational activation of ORs at discrete and additional internal membrane locations, including the somatic Golgi apparatus and Golgi outposts throughout the dendritic arbor, which are inaccessible to peptides and do not require receptor trafficking for activation.

Together, these results support a model in which OR activation occurs in living neurons through spatially (Figure 7A) and temporally (Figure 7B) resolved ‘waves’. Opioid peptides drive a ‘core’ activation pattern that is comprised of sequential waves of OR activation in the PM and propagating to endosomes. Agonist and antagonist drugs distort this pattern, with clinically relevant alkaloid agonists, such as morphine, uniquely driving a third wave of Golgi-localized OR activation in the soma and dendrites.

**Figure 7:**
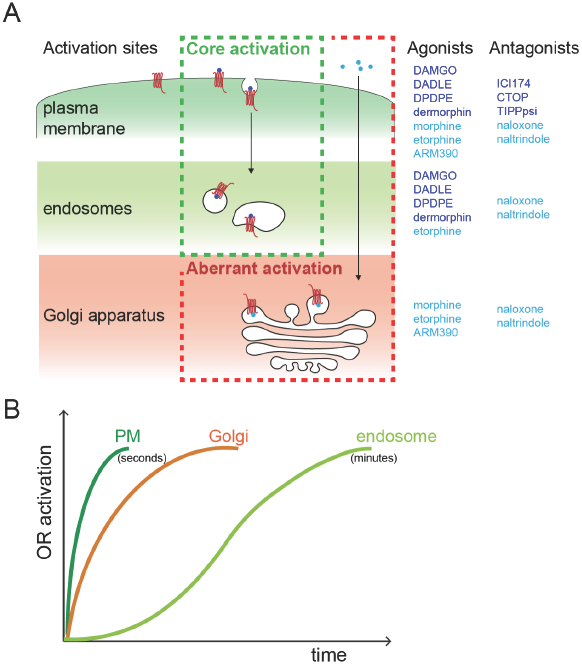
Spatiotemporal landscape of OR activation in the cell. **(A)** Summary of findings: Activation of OR occurs at distinct cellular membrane compartments in a ligand-dependent manner. Peptides agonists (dark blue) drive a ‘core’ activation pattern, with two sequential waves of OR activation, first in plasma membrane and then in endosomes following internalization of the receptor. Non-peptide agonist (light blue) distort this pattern by activating a Golgi-localized internal OR pool (‘aberrant’ activation). The distinct OR activation sites are differentially affected by peptide (dark blue) and non-peptide (light blue) antagonists. **(B)** Kinetics of OR activation at distinct cellular membrane compartments. Activation of ORs at PM and the Golgi apparatus occurs within seconds of agonist application. The endosome activation wave lags several minutes.

The present results have specific implications for opioid biology and therapeutics. Differences in the cellular actions of opioid drugs are presently understood to arise through ligand-selective bias imposed on the conformational landscape of individual receptors (Kenakin, 2011; Raehal et al., 2011; Staus et al., 2016). The present results identify an additional layer of selectivity imposed through ligand-specific bias of the spatiotemporal landscape of OR activation in individual neurons. An important future direction is to determine the functional significance of location-specific OR activation. A number of GPCRs are now recognized to initiate signaling from internal membranes, including endosomes and the Golgi apparatus (Calebiro et al., 2009; Irannejad et al., 2017; 2013; Vilardaga et al., 2014), and location-specific activation has been demonstrated to produce distinct downstream effects by affecting signal duration as well as pathway selectivity through proximity to downstream effectors (Godbole et al., 2017; Jong et al., 2014; Tsvetanova and Zastrow, 2014). Location-specific activation of ORs therefore suggests an expanded framework for understanding diversity and specificity of cellular opioid actions. In this regard, the Golgi-localized activation wave is of particular interest because it is robustly induced by opiate alkaloid drugs but not by peptide ligands.

The present findings also have more general implications for neural cell biology and neuropharmacology. Most neuromodulator receptors belong to the GPCR superfamily, and neurons represent some of the most structurally elaborate cells in the body. Thus, the spatiotemporal activation landscape that we delineate here for ORs may be widely applicable to neuromodulator receptors, particularly those whose natural ligands are hydrophilic. We also note that internal membrane pools of various GPCRs (including ORs) have been previously identified in neurons, but these were generally thought to represent inactive reserves for later mobilization to the PM. That such internal pools are subject to ligand-dependent activation could be broadly relevant to GPCR-directed drug action because most CNS-accessible drugs in current use are relatively hydrophobic, and thus are likely to be sufficiently membrane-permeant to effectively access internal membrane pools at the level of individual target neurons.

## Acknowledgements

We thank Seksiri Arttamangkul, Brian Kobilka, and John Williams for valuable discussion and providing critical reagents. We thank members of the Kobilka and von Zastrow laboratories for valuable discussion and advice. Some of the imaging experiments were carried out in the UCSF Nikon Imaging Center directed by DeLaine Larsen. We thank Simon Braun for advice and critical reading of the manuscript. This study was supported by research grants from the NIH (DA010711 and DA012864 to M.vZ., DA004443 to P.S., and DP5 OD 02304801 to A.M.) and the Canadian Institutes of Health Research (MOP-89716 to P.S.). M.S. is supported by the Swiss National Science Foundation (P2EZP3_152173 and P300PA_164712). D.J. received support from the UCSF Program in Breakthrough Biomedical Research.

## Author Contributions

M.S and M.vZ. conceived the experiments and wrote the manuscript. M.S. and D.J. performed the experiments and analyzed the data. A.M., T.L., J.S., and P.S. provided essential reagents and contributed to experimental design.

## Materials and Methods

### Cell culture, cDNA constructs, and transfection

HEK293 and HeLa cells (ATCC) were grown in DMEM supplemented with 10% FBS. Stably transfected HEK293 cells expressing N-terminally FLAG-tagged μ‐ or δ-OR (described previously) were cultured in the presence of 250 μg/ml Geneticin (Gibco). EGFP‐ and pmCherry-OR-sensor was created by amplifying Nb33 DNA using 5‘-3‘ TCGAAGCTTCCGGTAGCGGCAGCGGTATGGCACAGGTGCAGCTGG and TGCGGATCCTTATGCGGCCCCGTGATGGTG primers, adding the linker sequence GSGSG, and inserted into EGFP-C1 or pmApple-C1 vectors using HindIII and BamHI sites. GalT-mRFP1 (GalT aa 1-82) (Irannejad et al., 2017), EEA1-DsRed2 (EEA1 aa 1262-1411) and CLC-DsRed (Irannejad et al., 2013), and N-terminal signal sequence FLAG (ssf)-tagged murine μ-OR, δ-OR, and muscarinic acetylcholine receptor (M2) constructs in pcDNA3.1 vectors were previously described. For transient transfections, Lipofectamine 2000 (Invitrogen) was used according to manufacturer’s instructions. For live cell imaging, cells were plated on poly-L-lysine-coated 35-mm glass-bottomed culture dishes (MatTek Corporation) 48 h before the experiments. Cells were transfected 24h prior to imaging. Per culture dish, 200ng cDNA was used for OR-sensor or organelle markers (EEA1, GalT, CLC), and 1.5μg DNA was used for receptor constructs.

### Striatal neuron isolation, cDNA constructs, and transfection

Striatal neurons were prepared from embryonic day 18 Sprague-Dawley rats. The striatum, including the caudate-putamen and nucleus accumbens, was identified as described (Ventimiglia and Lindsay, 1998). Striata were dissected in ice cold Hank’s Buffered Saline solution, Calcium/magnesium/phenol red-free (UCSF Cell Culture Facility). Striata were dissociated in 1x trypsin/EDTA for 15 minutes at 37°C. Cells were washed in Gibco DMEM (Invitrogen) supplemented with 10% fetal bovine serum (UCSF, Cell Culture Facility) and 30 mM Hepes, then mechanically separated with a flame-polished Pasteur pipette. Striatal neurons were transfected using electroporation (Rat Neuron Nucleofector Kit; Lonza) before plating or using Lipofectamine 2000. Cells were plated on poly-D-lysine-coated 35mm glass-bottom dishes (MatTek) in DMEM (Invitrogen) supplemented with 10% FBS. Media was exchanged 1-4 days after plating for Neurobasal (Invitrogen) supplemented with Glutamax 1x (Invitrogen) and B27 1x (Thermo Fischer). Half of the media was exchanged every week, Cytosine arabinosine 2μM was added at 8 DIV. Transfection using Lipofectamine 2000 was performed on DIV 8, using 1μl of Lipofectamine and 1μg DNA in 1ml of media per 35mm imaging dish, media was exchanged 6 hours later. Cells were maintained in a humidified incubator with 5% CO2 at 37°C and imaged at 10-15 DIV. For transient neuronal expression, cDNA of ssf-μ-OR, ssf-δ-OR, EGFP-OR-sensor, ssf-μ-OR-EGFP, ManII(1-112)-pmApple was cloned into pCAGGS/SE expression vector with CAG promoter, using EcoR1/KpnI restriction sites.

### Reagents

DAMGO (E7384), DADLE (E7131), Carbamoylcholine chloride (Carbachol, C4382) morphine sulfate (1448005), Naloxone hydrochloride dehydrate (N7758), Forskolin (F6886), and M1 monoclonal FLAG antibody (F3040) were purchased from Sigma-Aldrich. AR-M 1000390 hydrochloride (4335), ICI 174,864 (0820), CTOP (1578), and Naltrindole hydrochloride (0740) were purchased from Tocris. Fluorescent Dermorphin (Arttamangkul et al., 2000) and TIPP(psi) (Schiller et al., 1993) have been previously been described. Alexa Fluor 647 and 555 Protein Labeling Kits (A20173, A20174) and Alexa Fluor-conjugated secondary antibodies, ProLong Gold antifade reagent were purchased from Invitrogen, anti-Giantin rabbit antibody (PRB-114C) from Covance. Drugs were added by perfusion or bath application at concentrations indicated in figure legends.

### Total internal reflection fluorescence microscopy

Live cell image series testing OR-sensor recruitment to the PM were acquired on a Nikon TE-2000 inverted microscope equipped with 60x 1.45 NA Plan Apo TIRF objective, a Bioptechs objective warmer and an Andor iXon EM+ EMCCD camera. The illumination from an argon laser source was focused on the periphery of the back focal plane of the objective with a micrometer guided illuminator (Nikon) to achieve total internal reflection. For perfusion, an insert was 3D printed and placed inside the imaging dish where it left a dead volume of about 300 ul. It was used to perfuse solution (Hepes buffered saline (HBS) with 135 mM NaCl, 5 mM KCl, 0.4 mM MgCl2,1.8 mM CaCl2, 20 mM Hepes, 1 mM d-glucose, and 2% FCS adjusted to pH 7.4 and 300–315 mOsmol/l) containing agonists or antagonists at indicated concentrations with a flow rate of 1.5ml/min.

Other TIR-FM imaging experiments were performed at 37 °C using a Nikon Ti-E microscope equipped for through-the-objective TIR-FM with a temperature‐, humidity‐ and CO2-controlled chamber (Okolab) and an Andor DU897 EMCCD camera. Images were obtained with a 100× 1.49 NA Apo TIRF objective (Nikon) with solid-state lasers of 405, 488, 561 and 647 nm (Keysight Technologies). Cells were imaged in DMEM without phenol red supplemented with 30 mM HEPES, pH 7.4 (UCSF Cell Culture Facility).

### Spinning disc confocal imaging

Live-cell spinning disc confocal imaging was carried out using Yokagawa CSU-22 confocal Nikon Ti microscope with a 60x or 100x 1.4 NA Plan Apo VC oil objective and a temperature-, humidity‐ and CO2-controlled Okolab enclosure Bold Line incubator. 488-nm, 568-nm and 640-nm Coherent OBIS lasers were used as light sources. Time-lapse images were acquired with an Evolve Delta EMCCD camera (Photometrics) driven by Micro-Manager 1.4 or Nikon Elements 4.5. Cells were imaged every 2-15 sec for up to 25 min in DMEM without phenol red supplemented with 30 mM HEPES, pH 7.4 (UCSF Cell Culture Facility). Agonists or antagonists were added by bath application during acquisition. For immunofluorescence imaging, cells were cells grown on coverslips and fixed using 4% formaldehyde in PBS. Cells were permeabilized with 0.05% saponin and 1% BSA in PBS and incubated with primary (1:1000) and secondary (1:1000) antibodies. Specimens were mounted using ProLong Gold.

### Fluorescence recovery after photobleaching

FRAP experiments of EGFP-OR-sensor at endosomes were performed at 37 °C on a Andor Borealis CSW-W1 spinning disk confocal Nikon Ti Microscope with Andor 4-line laser launch and a temperature-, humidity‐ and CO2-controlled chamber (Okolab). HEK293 cells transiently expressing EGFP-OR-sensor and stably expressing ssfMOR (labeled with M1 anti-Flag labeled with Alexa 647) were imaged using a 100× 1.49 NA Apo TIRF objective (Nikon) and an Andor Zyla 4.2 sCMOS camera controlled by MicroManager software. Cells were incubated with 10 μM DAMGO for 15 min prior to imaging. EGFP-OR-sensor was bleached at 2-3 individual receptor-loaded endosomes per cell using a Rapp Optoelectronic UGA-40 photobleaching system with 473 laser (Vortran). EGFP fluorescence recovery was monitored by acquiring images every 2 seconds for 5 min. The receptor signal was acquired throughout the image series to correct for possible movement of endosomes. The mean EGFP fluorescence intensity at bleached endosomes at each time point was corrected for background signal and photobleaching of the cell. Fluorescence intensity before bleaching was normalized to 1 and directly after bleaching to 0.

### Quantitative image analysis

All quantitative image analysis was performed on unprocessed images using MatLab or ImageJ. For quantifying the enrichment of OR and OR-sensor at endosomes, Golgi apparatus, or clathrin-coated pits, time-lapse movies were divided in sub-regions containing the fluorescent marker proteins used for labeling (EEA1 (aa 1262-1411), GalT (aa 1-82), or CLC). Movies were thresholded to encompass signal from marker proteins throughout the movie and to generate a mask of the structures. Filtering to exclude regions smaller than 5 pixels was applied, remaining objects were smoothened with a dilatation of 1 pixel before an erosion of 1 pixel. The generated mask was applied to the OR or OR-sensor channel to measure the fluorescence at the location of marker objects. The average fluorescence in the mask was background subtracted and normalized to baseline (before agonist addition) or to the cellular fluorescence outside the thresholded structures. To determine the linear correlation coefficient between agonist-loaded endosomes (marked by dermorphin) or Golgi outposts (marked by ManII) and OR-sensor or soluble EGFP, a 50-pixel line was drawn along the dendrite on individual endosomes identified in the marker channel. The fluorescence profile obtained in both channels for each endosome was normalized with the maximal value set to 1 and the minimal value set to 0. Linescans were aligned to the maximal value in the red channel and the linear correlation coefficient calculated for every endosome as the value of the GFP channel for this linescan point.

### Luminescence-based cAMP assay

Cellular cAMP was detected in living cells as described previously (Irannejad et al., 2013). In brief, HEK293 cells, stably expressing ssf-μ-OR were transfected with GloSensor-20F cAMP reporter (Promega), and EGFP-OR-sensor or empty EGFP-C1 vector and plated in 24-well dishes. 24 hours post transfection, cells were treated with Forskolin and varying concentrations of DAMGO. Images were collected every 10 s for 30 min using a CCD (Hamamatsu C9100-13) in a light-proof cabinet at 37 °C. Maximum luminescence values were measured using ImageJ and normalized to the maximum luminescence measured in the presence of 2 μM Forskolin.

